# Regulation of oogenesis in the queen honey bee (*Apis mellifera*)

**DOI:** 10.1101/2021.03.08.434480

**Authors:** Sarah E Aamidor, Carlos AM Cardoso-Júnior, Januar Harianto, Cameron J Nowell, Louise Cole, Benjamin P Oldroyd, Isobel Ronai

**Affiliations:** Behaviour and Genetics of Social Insects Laboratory, Ecology and Evolution, School of Life and Environmental Science, Macleay Building A12, University of Sydney, NSW, 2006, Australia; Departamento de Biologia Celulare Bioagentes Patogênicos, Faculdade de Medicina de Ribeirao Preto, Universidade de Sao Paulo, Ribeirao Preto, Brazil; School of Life and Environmental Science, Macleay Building A12, University of Sydney, NSW, 2006, Australia; Drug Discovery Biology, Monash Institute of Pharmaceutical Sciences, Monash University, 381 Royal Parade, Parkville 3052, Victoria, Australia; Microbial Imaging Facility, I3 Institute, Faculty of Science, The University of Technology Sydney

**Keywords:** ovary activation, ovariole, nurse cell chamber, honeybee, eusociality, phenotypic plasticity

## Abstract

In the honey bee (*Apis mellifera*), queen and worker castes originate from identical genetic templates but develop into different phenotypes. Queens lay up to 2,000 eggs daily whereas workers are sterile in the queen’s presence. Periodically queens stop laying; during swarming, when resources are scarce in winter and when they are confined to a cage by beekeepers. We used confocal microscopy and gene expression assays to investigate the control of oogenesis in honey bee queen ovaries. We show that queens use different combination of ‘checkpoints’ to regulate oogenesis compared to honey bee workers and other insect species. However, both queen and worker castes use the same programmed cell death pathways to terminate oocyte development at their caste-specific checkpoints. Our results also suggest that the termination of oogenesis in queens is driven by nutritional stress. Thus, queens may regulate oogenesis via the same regulatory pathways that were utilised by ancestral solitary species but have adjusted physiological checkpoints to suit their highly-derived life history.

**Summary statement:** Honey bee queens regulate oogenesis using a different combination of ‘checkpoints’ to workers, but both castes use the same molecular pathways.

## Introduction

In eusocial insects, females develop into either a highly reproductive queen or a non-reproductive worker (Oster and Wilson 1978). In the honey bee, *Apis mellifera*, queens and workers arise from the same genetic template (genotype), but have radically different morphological and behavioural phenotypes (Snodgrass 1956; Dade 1977; Winston 1987). Queens lack worker-specific characters such as a barbed sting, pollen baskets (corbiculae) and a honey crop, whereas workers generally lack the specialised reproductive organs of the queen such as the sperm-storing spermatheca. Notably, while both queens and workers have ovaries, they differ enormously in the number of ovarioles (the tubes in which oocytes develop). Queens have 150-200 ovarioles per ovary, whereas workers have approximately four (Snodgrass 1956; Rozen 1986; Ronai, Allsopp et al. 2017). The queen’s ovaries occupy the majority of her abdominal cavity and can produce up to 2,000 eggs per day (Page and Erickson 1988). Honey bee queens therefore have one of the highest levels of fecundity of any insect (Hodin 2009).

In an activated honey bee ovary, each ovariole contains multiple oocytes (immature eggs) at different stages of development. Oogenesis begins at the tip of the ovariole when a germline cell divides repeatedly to produce what will become an oocyte and a cluster of accompanying nurse cells. The nurse cells provide nutrition and other elements required by the developing oocyte as it travels through the ovariole, in a conveyor-belt fashion, towards the median oviduct (Snodgrass 1956). As the oocyte approaches the oviduct the nurse cells transfer their contents to the oocyte in a process known as nurse cell ‘dumping’ (Spradling 1993; Cavaliere, Taddei et al. 1998), and die as the oocyte completes its development. Finally, the mature oocyte passes through the median oviduct and is laid.

Worker honey bees, on the other hand, are facultatively sterile (Visscher 1989; Ratnieks 1993). Young workers can switch between ‘deactivated’ ovaries when the queen is present to ‘activated’ when the queen is absent (Malka, Shnieor et al. 2007). The mechanisms underlying the reproductive plasticity of workers has been widely investigated (e.g Hoover, Keeling et al. 2003; Malka, Shnieor et al. 2007; Woyciechowski and Kuszewska 2012; Traynor, Le Conte et al. 2014; Ronai, Oldroyd et al. 2016; Ronai, Oldroyd et al. 2016). Young workers continuously produce new oocytes, but in the presence of a queen, the developing oocytes are aborted during mid-oogenesis via programmed cell death (PCD) (Tanaka and Hartfelder 2004; Tanaka, Schmidt-Capella et al. 2006; Ronai, Barton et al. 2015). The reproductive plasticity of young workers allows them to rapidly activate their ovaries, by switching off the PCD mechanisms, thereby allowing pre-existing immature oocytes to develop. This mechanism likely evolved because there is a fitness premium for the first workers to successfully lay eggs in a queen-less colony (Page and Robinson 1994). A queen signals her presence to the workers via pheromones (Butler 1957; Butler and Fairey 1963; Hoover, Keeling et al. 2003), the most significant of which is queen mandibular pheromone (Butler, Callow et al. 1961). In a colony with a queen, the presence of queen mandibular pheromone and brood pheromones in a worker’s environment activates the gene network(s) that initiate PCD in early oocytes of workers (Le Conte, Mohammedi et al. 2001; Ronai, Oldroyd et al. 2016; Hartfelder, Tiberio et al. 2018). In contrast to workers, the mechanisms underlying the reproductive plasticity of mated queens is unlikely to be regulated by queen mandibular pheromone and brood pheromones, for if they were influenced by these cues, they would be permanently sterile. Nonetheless, queens, like workers, are able to curtail oogenesis in response to environmental cues.

Mated queens exhibit reproductive plasticity under three specific circumstances. First, in temperate zones, queens drastically reduce egg laying in winter (Seeley 1985), becoming reproductively active again in late winter or early spring (Avitabile 1978; Shehata, Townsend et al. 1981). This annual plasticity in oogenesis is likely triggered by decreased availability of fresh pollen combined with declining day length (Mattila and Otis 2007). Second, reproductive swarming; when the queen and approximately half of the colonies workers depart the nest to establish a new colony (Simpson 1959). In preparation for swarming, the workers reduce the frequency at which they feed the queen (Allen 1956; Pierce, Lewis et al. 2007). Probably as a consequence of this reduced feeding, the queen’s egg production declines significantly (Allen 1956; Grozinger, Richards et al. 2014). As a result the queen’s weight is significantly reduced, which likely allows her to fly more easily when she leaves with the swarm (Seeley and Fell 1981). After the swarm leaves the natal nest, the bees hang in a cluster nearby. While in the cluster the queen’s egg production almost ceases (Pierce, Lewis et al. 2007). Only once the swarm moves to its new home and new brood cells are built, does the queen resume egg laying (Ambrose 1976; Winston 1987). Thirdly, commercial queens used in agriculture are often forced to undergo periods when they are prevented from laying due to beekeeping practises (Rhodes, Somerville et al. 2004). Queens produced for sale are sometimes ‘banked’ within individual cages in a colony for several weeks (Laidlaw and Page 1997). Banked queens are cared for by the colony’s workers but lack a comb in which to lay eggs. Queens are often shipped via mail in individual cages with an escort of 5-8 workers that feed and groom the queen, usually for a period of up to 10 days (Laidlaw and Page 1997). Caged queens do not lay eggs, have restricted food, and their abdomens become much reduced in size (Laidlaw and Page 1997). Nonetheless, once the queen is introduced to her new colony she recommences egg laying in a few days (Rhodes, Somerville et al. 2004).

In this study, we investigate the mechanisms that enable honey bee queens to switch between activated and deactivated ovaries. We studied oogenesis in queens by examining the morphological changes in their ovaries when they are prevented from laying in two different social environments; (i) within the colony and (ii) when removed from the colony with a few escorting workers. Since queens and workers originate from the same genetic template, we further investigated whether the reproductive plasticity of queens is regulated by differential expression of the same genes that are associated with the reproductive plasticity of workers, and whether such plasticity is driven by nutritional stress. Specifically, we quantified the expression of genes associated with: PCD (*Anarchy, Buffy, Apaf-related killer* gene [*Ark*], *long non-coding ovary-1 [Lncov1], Tudor-SN*, and *Draper*); nutrition (target of rapamycin [Tor], *Forkhead box O* [*FoxO*], elongation growth factor receptor [*Egfr*]); Juvenile Hormone production (*Krüppel-homolog 1* [*Kr*-*h1*]), and vitellogenesis (Vitellogenin [Vg] and Vitellogenin Receptor [*VgR*]). We hypothesise that the genetic pathways controlling oogenesis in queens would be similar to those seen in workers (i.e. associated with PCD). We further hypothesised that in contrast to workers, where queen mandibular pheromone and brood pheromones are the cue for workers to stop oogenesis, in queens it is nutritional cues that regulate oogenesis. More broadly, we note that in solitary Hymenopterans, nutritional cues are intimately related to oogenesis (Toth, Varala et al. 2007). Thus nutritional cues may also have a role in regulating oogenesis in the queens of advanced eusocial insects (Hunt and Amdam 2005; Dolezal, Flores et al. 2013).

## Methods

### Biological material

Queens (n = 25) were removed from their colonies during summer (December 2018). All individuals were mature (> 6 weeks old), mated, laying queens of standard Australian commercial stock mostly derived from *A. m. ligustica*.

Queens were randomly assigned to three treatment groups. First, fifteen queens were placed in individual plastic mailing-introduction cages (4 cm x 2 cm, JZ-BZ Queen Cages) used by commercial beekeepers, with five to eight supporting workers and kept in a dark at room temperature. The workers had ad *libitum* access to water and queen candy (a mixture of caster sugar and honey) to feed the queens. Collectively we refer to these treatments as caged ‘ex-colony’ (EC). Five queens from this EC group were collected on days 4, 7 and 10 after caging (which is within standard beekeeping practice). Second, five queens with no escorting workers were suspended in their cages between brood frames above a queen excluder in a strong colony. The queen excluder prevented access by the resident queen to the caged queens. Queens were fed by host workers through the mesh of their cages. These queens were collected 10 days after caging. We refer to these queens as caged ‘in colony’ (IC). Third, the remaining five queens were collected from their colonies at day 0 as the control group. During the course of these experiments two queens died, one in the 10-day EC group and one in the IC group. Hence there is no data from these two queens.

All collected queens were immediately frozen on dry ice and stored at −80 °C. Queens were dissected in RNase-free phosphate-buffered saline (PBS) and their paired ovaries were delicately removed. Both ovaries were imaged under a dissecting light microscope (Olympus SZ40 Stereo Zoom) at 8X magnification in order to measure the ovary 2D area. One of the paired ovaries was randomly chosen and halved. One half was set aside for confocal microscopy and the other for gene expression analysis. The remaining ovary was stored but not used for this study. All dissected ovaries were stored at −80 °C until required.

### Gene expression in the ovaries

Total RNA was extracted from each ovary sample (n = 23) using TRIzol (Invitrogen). The amount of RNA present per sample was estimated using a Qubit 2.0 Fluorometer (Invitrogen) and samples were diluted to 200 ng/µL with RNase-free water. DNA was removed using a Turbo DNase Kit (Thermo Fisher Scientific). For each sample, cDNA was synthesised from 320 ng of RNA (64 ng/µL) using a SuperScript™ III Reverse Transcriptase Kit (Invitrogen) with Oligo (dT) universal primers with appropriate positive and negative controls. All cDNA samples were diluted to 5 ng/µL with RNase-free water.

We quantified the expression of 11 genes of interest (*Anarchy, Buffy, Draper, Egfr, Lncov1, Tudor, Tor, FoxO, Kr-h1, Vg and VgR*) in the queen ovaries using RT-qPCR. Each reaction was run in triplicate with a final volume of 5 µL [1 µL cDNA (5 ng/µL), 2.5 µL SYBR Green (Applied Biosystems, Life Technologies) and 1.25 pmol of each primer] using a CFX384 Real-Time System (Bio-Rad). The sequences of the primers and reaction conditions for the 11 genes of interest and four candidate reference genes (*pRS5, RP49, Ef1a* and *Actin*) are provided in Supplementary Table S1. The reaction efficiency for each primer set was obtained by a standard curve (serial dilution) fitted for each of the genes (Supplementary Table S1). Of the four candidate reference genes, *Ef1a* and *Rf49* were found to have the most stable expression using the software BestKeeper (Pfaffl, Tichopad et al. 2004). These two genes have been previously recommended as appropriate reference genes for gene expression studies in honey bees (Lourenço, Mackert et al. 2008; Reim, Thamm et al. 2013). Relative gene expression was calculated according to the efficiency correction formula considering the efficiency of each gene and normalised with the two reference genes (Brito, McHale et al. 2010).

### Fixing, staining and confocal imaging of the ovaries

Ovaries were prepared for staining according to Dallacqua and Bitondi (2014). Briefly, each ovary was fixed and permeabilised in heptane fixative (0.83% heptane, 0.067% HEM buffer (pH 6.9, 0.1M HEPES, 2mM MgSO4 and 1mM EGTA), 0.01% paraformaldehyde and 0.0167% DMSO) for 30 minutes on a shaker, followed by three washes in PBS. The ovaries were then dehydrated with increasing concentrations of ethanol (25%, 50%, 75% and 100%) and stored over night at −20 °C. For staining, the ovaries were rehydrated with decreasing concentrations of ethanol in water (75%, 50% and 25%) and then washed three times in PTw (PBS and 0.1% Tween20). Ovaries were refixed in Triton fixative (PBS, 0.1% Triton X-100, 4% paraformaldehyde and 10% DMSO) for 20 min and washed three times in PTw. In order to further permeabilise the tissue, samples were incubated for 10 minutes in 20 µg/mL proteinase K followed by two washes in 1% glycine and three washes with PTw. The ovaries were refixed in Triton fixative for 20 minutes and washed six times in PTw to remove all traces of fixative. Ovaries were then stained for actin with ActinGreen 488 (Invitrogen) for 30 minutes, washed three times with PTw and counterstained for nucleic acid with DAPI (0.004% v/v) (Sigma) for five minutes followed by three washes in PTw. The ovaries were then split into clusters of one to four ovarioles and mounted on slides in Fluoroshield (Sigma) with glass cover slips. (During fixation one ovary from the control group was lost and therefore not imaged). Slides were imaged using a ZEISS LSM 800 confocal microscope using 450 and 488nm excitation wavelengths to image the nucleic acid and actin respectively. Single focal plane and z-stacks were taken using either a 10x or 20x objective. Z-stack images were processed to create a two-dimensional maximum intensity projection (MIP). All images were acquired, stitched and processed using ZEN blue software.

### Image analysis

Images were analysed using a custom macro in the Fiji distribution of ImageJ (Schindelin *et. al*., 2012). Stereo images were used to calculate total area of both ovaries (one image for each individual ovary). Confocal images were enhanced using the ImageJ gamma function at 0.6 to increase contrast. For each stitched image, the oocytes and Nurse Cell Chambers (NCC) in each ovariole were traced by hand and the resulting ROIs saved for future validation if required.

Confocal image analysis is described in detail in Supplementary Materials and Methods S1 and available for download from https://doi.org/10.26180/13557764. Briefly, oocytes and NCC in each string were outlined using the freehand region of interest (ROI) tool on maximum Z projections of each image set from the most developed intact oocyte (oviduct end) until the oocytes and NCC could no longer be distinguished in the late germarium (Büning 1994). Each ROI was then used to calculate the equivalent spherical radius by using the maximum and minimum Feret’s diameter to first calculate the oblate volume (due to the ‘objects’ - oocyte or NCC - being non-spherical) of the object and then to calculate the equivalent spherical radius. The data was logged and then further analysed using Excel and GraphPad Prism. In order to visualise the changes along the entire ovariole a locally weighted smoothing (LOESS) function was added on the scatterplot output using the ‘loess()’ function from the base ‘stats’ package in R (version 4.0.3).

### Statistical analysis

To determine the effects of treatment group on the morphological features and gene expression of the ovariole, we modelled the data using generalised linear models (GLM) with a normal link function. The dependent variable was unit size or position in the ovariole, or the fold change in gene expression. The independent (fixed) factor was the treatment group (control (Day 0), EC day 4, EC day 7, EC day 10 and IC day 10).

All covariates were ordinal, thus model significance was tested with an analysis of variance (ANOVA) using the Wald Chi-square test statistic and Type 3 error. Where statistical results indicated significant differences at p < 0.05, estimated marginal means (EMM) we used to develop linear estimates of pairwise comparisons and then tested using Tukey’s honestly significantly different (HSD) test to identify significant differences at p < 0.05.

Analyses were performed in R (version 4.0.3). GLMs were performed using the ‘Anova()’ function from the ‘cars’ package (version 3.0-10) while post-hoc analyses were performed using the ‘emmeans()’ function from the ‘emmeans’ package (version 1.5.3).

## Results

### Oogenesis in a laying queen

Below we provide a description of normal oogenesis, obtained from our high-resolution confocal images of the ovaries of queens freshly taken from their colonies (controls; Day 0). Descriptions are also based on the existing honey bee ovary morphology literature (Snodgrass 1956; Gutzeit, Zissler et al. 1993; Büning 1994).

At the apical tip of each ovariole is the terminal filament (Figure 1 Bi) and germline stem cells (Figure 1 B). The early germarium stage is characterised by a single germ cell dividing repeatedly to produce a cluster of cells (also known as cystocyte rosettes) around actin polyfusomes (Figure 1 Bi). Subsequently, one of the cells differentiates into the oocyte, while the remaining cells of the cluster become the nurse cells that will nourish the developing oocyte. Actin fusomes also rearrange into ring canals (Figure 1 Bii), connecting the oocyte to its accompanying nurse cells.

**Figure 1.**
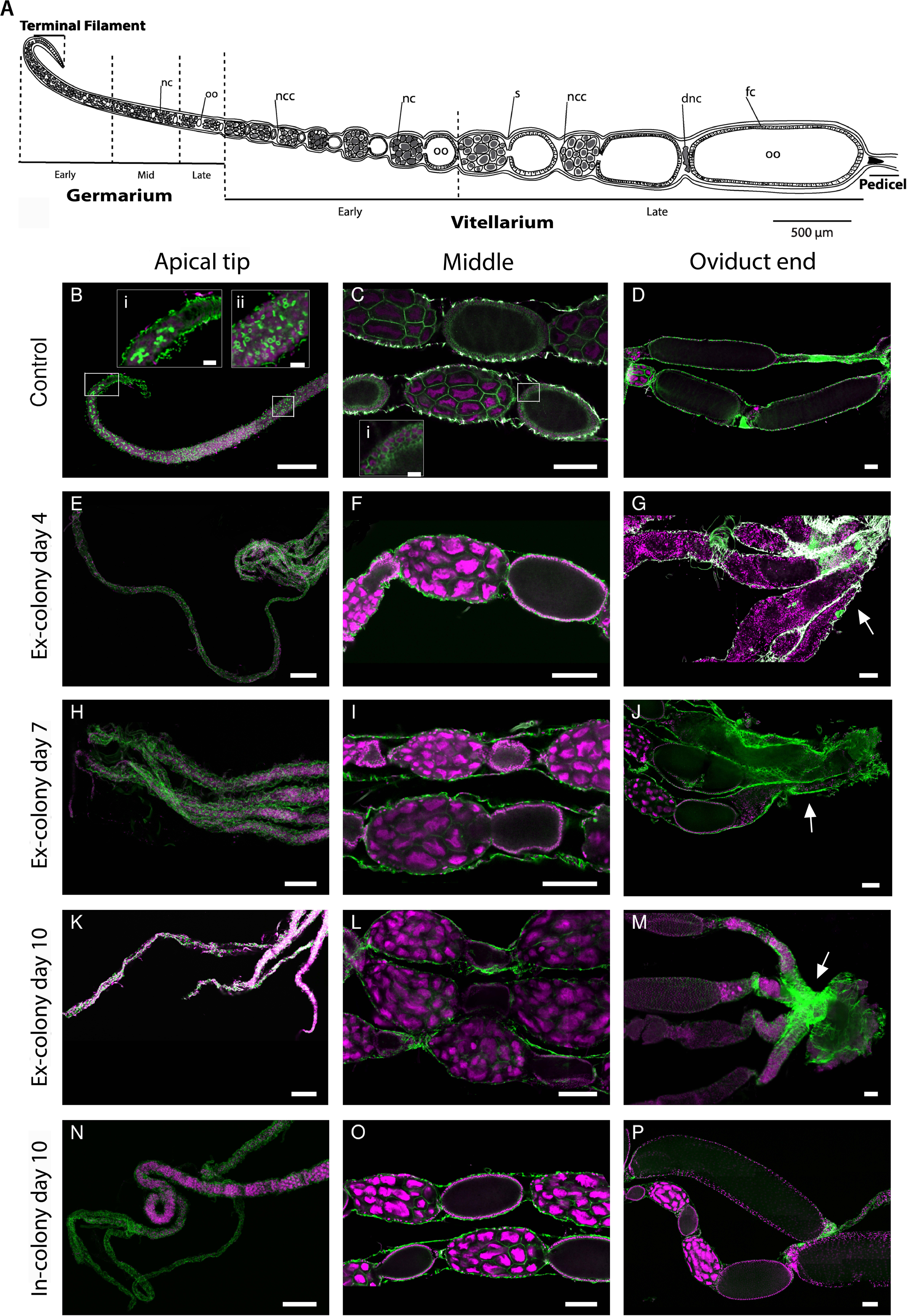
Oogenesis in honey bee queens. (A) A schematic representation of oogenesis in a mated laying queen with the four regions: terminal filament; germarium; vitellarium; and pedicel (nc = nurse cell, oo= oocyte, ncc = nurse cell chamber, s = sheath, dnc = dumped nurse cell, and fc = follicle cell); B-D. Ovaries of a control queen: (B) terminal filament and early germarium; (Bi) magnification of terminal filament and actin polyfusomes; (Bii) magnification of ring canals; (C) early vitellarium; (Ci) magnification of follicle cells; and (D) late vitellarium. E-G. Ovaries of a queen that had been caged EC for four days: (E) terminal filament and early germarium; (F) early vitellarium; and (G) late vitellarium showing actin aggregation (arrow). H-J. Ovaries of a queen that had been caged EC for seven days: (H) terminal filament and early germarium; (I) early vitellarium; and (J) late vitellarium showing actin aggregation (arrow). K-M. Ovaries of a queen that had been caged EC for ten days: (K) terminal filament and early germarium; (L) early vitellarium; and (M) late vitellarium showing actin aggregation (arrow). N-P. Ovaries of a queen that had been caged IC for ten days; (N) terminal filament and early germarium; (O) early vitellarium; and (P) late vitellarium. Ovaries were stained for nuclei (magenta) and actin (green). Scale bars 100 μm and magnified insets 10 μm.

During the mid-germarium stage, oocytes slowly increase in size and are distinguishable from their accompanying nurse cells (Figure 1 A). By the late germarium stage, the oocyte is located towards the oviduct end relative to its accompanying nurse cells (Figure 1 A).

The vitellarium stage is defined by the oocyte becoming encased in a layer of somatic follicle cells (Figure 1 Ci) and the accompanying nurse cells forming a Nurse Cell Chamber (NCC) (Figure 1 A and C). By both qualitative and quantitative analysis of the confocal images, we observed the oocyte and the NCC growing at different rates during the vitellarium stage (Figure 2). Initially the NCC is larger than the oocyte (Fig. 1C) until a point that we define as the ‘tipping point’ when the developing oocyte first exceeds the size of its accompanying NCC (Figure 1 A and 2). After the tipping point the late vitellarium stage begins and the oocyte grows rapidly (Figure 1A and 2). And the nurse cells undergo developmental programmed cell death (King 1970), ultimately transferring their remaining cytoplasm into the oocyte in a process termed nurse cell dumping (Figure 1 A and D, Figure 2). This nurse cell dumping results in the complete collapse of the NCC (‘dnc’ in Figure 1A; see also Figure 1D). At the final stage of oogenesis, the oocyte is fully formed and situated at the oviduct end of the ovariole (Figure 1A and 1D). The pedicel (also known as the basal stalk), a constriction at the entry to the oviduct which prevents the oocyte from moving into the oviduct until it is ready to be laid can also be seen clearly by confocal microscopy (Figure 1A and 1D).

**Figure 2.**
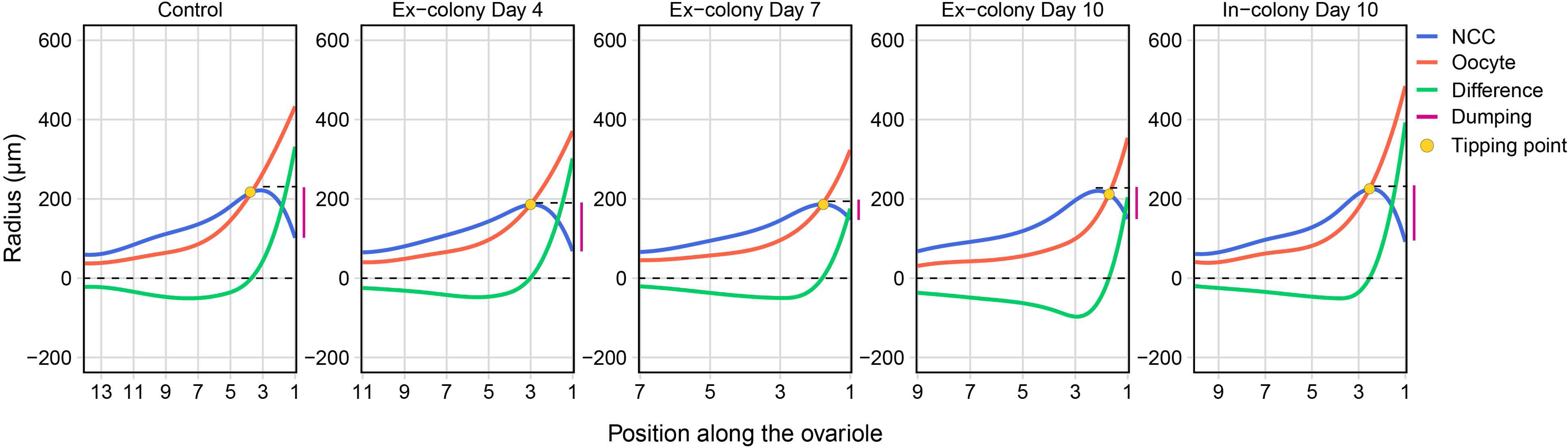
Dynamics of oogenesis in honey bee queens. The mean radius size (μm) of the oocytes (red line) and nurse cell chambers (NCC) (blue line) derived from averaged data for the control, caged EC days 4, 7 and 10 and caged IC for day 10 IC treatment groups. The difference between the size of the paired NCC and oocyte is indicated (green line). The cell position represents the unit number along the chain in the ovariole with the proximal oocyte at the oviduct end designated as position 0 (see Materials and Methods S1). The amount of NCC dumping (purple line) and tipping point (orange circle) are indicated for each group.

### Oogenesis in queens prevented from oviposition

Stereomicroscopy showed that caging, both ex-colony (EC) and in-colony (IC), resulted in abnormal ovaries. Overall, the longer a queen was caged EC, the smaller the ovaries became (Figure 3 A). We found that mean ovary area differed significantly among treatments (Wald χ^2^ = 96.25, d.f. = 4, *P* < 0.0001; Figure 3 B). Queens caged EC had significantly smaller ovary area compared to the controls. Although queens caged IC for 10 days did not show such an extreme reduction in ovary area, they too were smaller than the controls (Figure 3 B).

**Figure 3.**
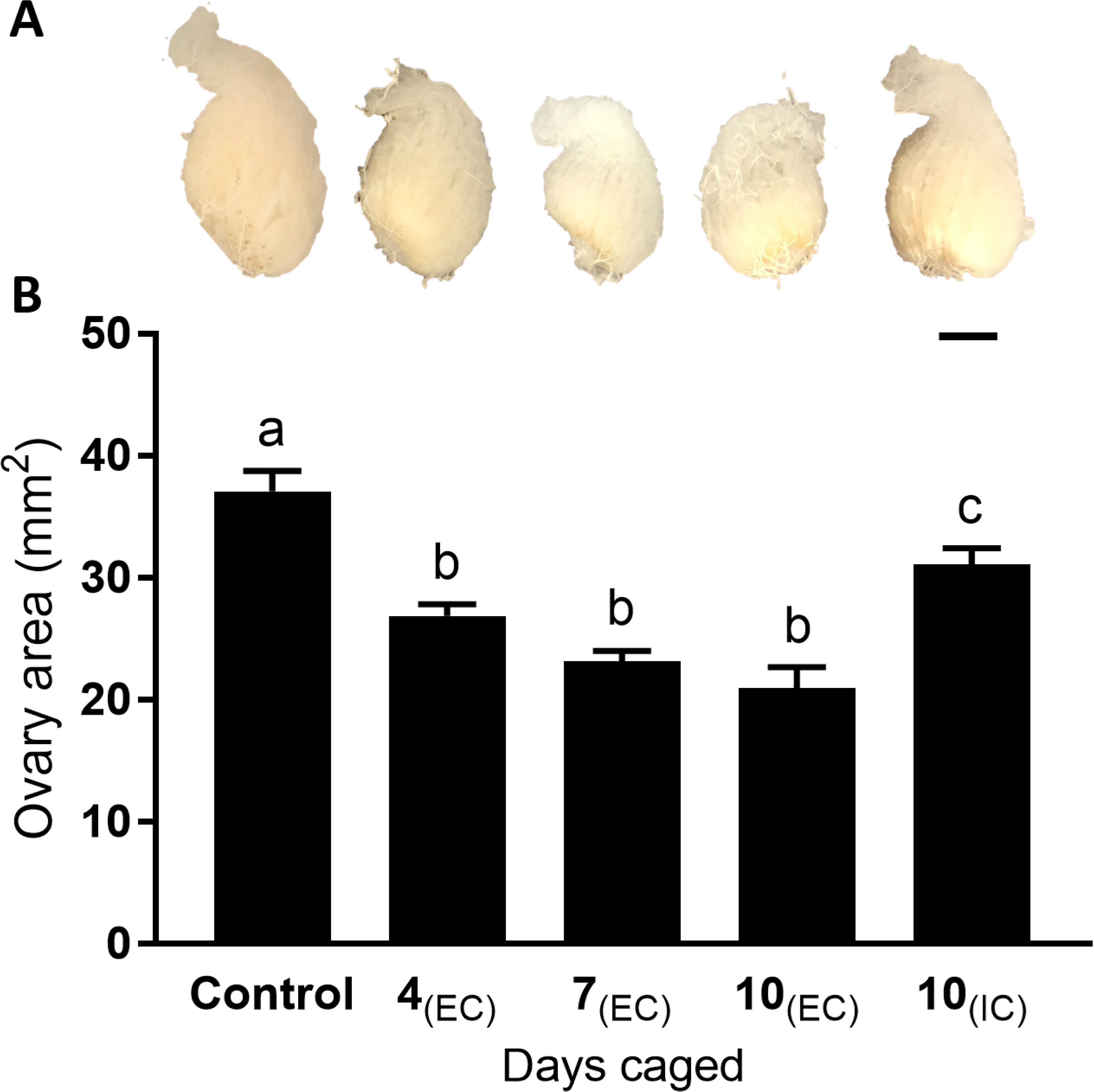
Change in ovary area of honey bee queens from the control, caged in-colony (IC) (4, 7 or 10 days) or caged Ex-colony (EC) (10 days) treatment groups. (A) Light microscopy image of a representative queen ovary from each treatment group (scale bar 2 mm). (B) Mean area (mm^2^) of queen ovaries across treatment groups (n = 4 control, caged IC 10 days and caged EC 10 days; n = 5 caged EC 4 and 7 days). Error bars are SE of the means. Different letters (a-c) represent statistical significance (*p* < 0.05).

Overall, confocal microscopy showed that there was no significant effect of caging treatments on the total number of oocytes (Wald χ^2^ = 6.04, *d*.*f*. = 4, *P* = 0.196; Figure S1). However, the size of the most mature (i.e. maximum-sized) oocyte was significantly reduced by the caging treatments (Wald χ^2^ = 18.79, *d*.*f*. = 4, *P* < 0.001; Figure 2 and S2 A). The size of the most mature oocyte was similar in queens that were caged EC for 10 days and the control queens (Figure S2 A). In contrast, the maximum size of the NCC was not significantly affected by the caging treatments (Wald χ^2^ = 8.97, *d*.*f*. = 4, *P* = 0.062; Figure 2 and S2 B). In addition, the minimum size of oocytes and NCC did not significantly change as a result of caging treatments (Wald χ^2^ = 5.43, *d*.*f*. = 4, *p* = 0.246; Wald χ^2^ = 1.48, *d*.*f*. = 4, p = 0.830; Figures S2 C and D respectively).

Although the number of identifiable oocytes per ovariole was unaffected by treatment (Figure S1), we nonetheless observed changes in the developmental gradient of oocytes along the ovarioles between treatments. The cells of the early germarium did not extend to the apical end of the ovariole in the caged queens (Figure 1 E, H, K and N). The early vitellarium stage in the EC caged queens had seemingly healthy NCCs, but the oocytes were misshapen and starting to deteriorate (Figure 4 A, arrows). This deterioration of the oocyte progressively intensified as the days of EC caging increased (Figure 1 I, L and 4 B, arrow). In contrast, the oocytes in the early vitellarium stage in IC caged queens (Figure 1 O) seemed similar to the control queens. The difference between the volume of the oocyte and its accompanying NCC provides a measure of the combined changes in the units (oocytes and NCC) along the ovariole (Figure 2, green line). In all the caged treatments the tipping point, marking the start of the late vitellarium stage, significantly shifted towards the oviduct end of the ovariole relative to control queens (Wald χ^2^ = 119.43, *d.f.* = 4, p = 0.0001; Figure 2 and S3). This shift suggests that there was a delay in the maturation of the oocytes in the caged queens, which in turn reduced the number of mature oocytes present in the ovariole. The tipping point for the IC caged queens was at a similar position in the ovariole to those seen in queens that had been caged EC for four days (Figure 2 and Figure S3).

**Figure 4.**
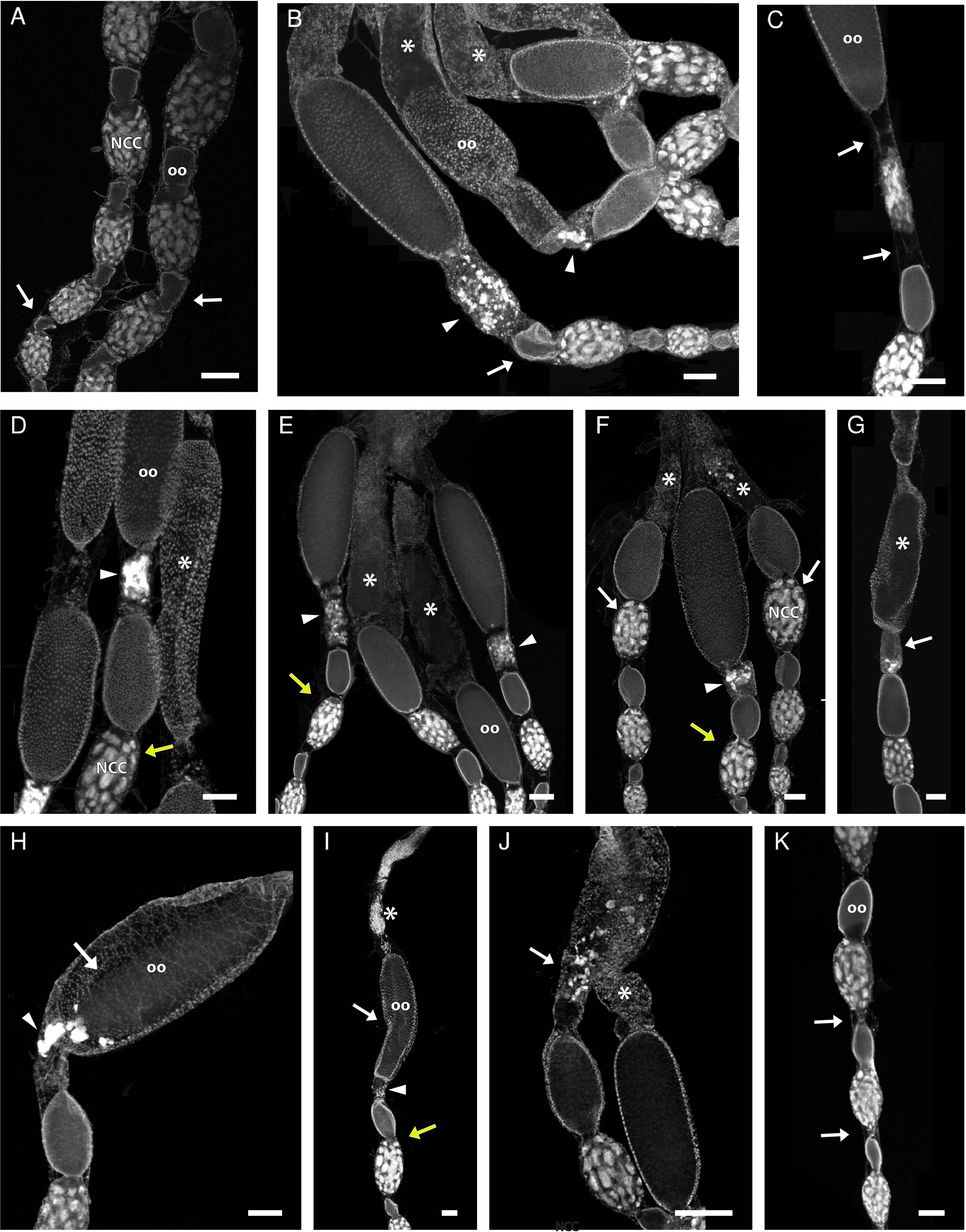
Oogenesis abnormalities in honey bee queen ovarioles after caging for 4, 7 and 10 days ex-colony (EC) and 10 days in-colony (IC). (A) EC four days early vitellarium. Misshapen oocytes (arrows) among healthy-looking nurse cell chambers; (B) EC 10 days late vitellarium. Some oocytes are underdeveloped and misshapen (arrow). There is also brightly-stained fragmented nucleic acid, suggesting premature programmed cell death of nurse cells but with no chamber collapse (arrow heads). Mature oocytes undergoing degradation (asterisk); (C) EC 7 days late vitellarium. The cytoplasm bridge is lost, disconnecting the oocyte from its nurse chamber and the connection between the oocyte and the next NCC is deteriorated (arrows); (D) EC 4 days late vitellarium. The nurse cell dumping (arrow head) is premature, occurring for an insufficiently-developed oocyte immediately after the ‘tipping point’ (yellow arrow). A mature oocyte is collapsing (asterisk); (E) EC 7 days early and late vitellarium. The nurse cell dumping (arrow heads) is premature, occurring immediately after the tipping point (yellow arrow). Mature oocytes are undergoing degradation (asterisk); (F) EC 10 days vitellarium. Premature nurse cell dumping (arrow head) is occurring immediately after the tipping point (yellow arrow) or no nurse cell dumping at all (arrows). The most developed oocytes are degrading (asterisk); (G) EC 4 days late vitellarium. Programmed cell death of the nurse cells has occurred, but the nurse cell chamber has not collapsed (arrow). A mature oocyte is collapsing (asterisk); (H) EC 7 days late vitellarium. The oocyte has disconnected from its surrounding follicle cells (arrow) indicating the beginning of its degradation. Brightly-stained fragmented nucleic acid are being transferred to the oocyte following NCC dumping (arrow head); (I) EC 10 days late vitellarium. Structural collapse of an oocyte (arrow) with premature NCC dumping occurring (arrow head) immediately after the tipping point (yellow arrow). The most mature oocyte is completely degraded (asterisk); (J) EC 10 days late vitellarium. The bright white dots in the mature oocyte are a sign of programmed cell death (arrow) and the most mature oocyte has completely collapsed (asterisk); (K) IC 10 days early vitellarium. The connection between the oocyte and its preceding NCC is deteriorating (arrows). oo = oocyte; NCC = nurse cell chamber. Ovarioles have been stained for nucleic acid (DAPI). Scale bars 100μm.

By the late vitellarium stage several structural abnormalities could be identified as a result of caging. In both EC and IC the cytoplasm bridges (ring canals) that normally connect the oocyte to its accompanying NCC were sometimes lost, resulting in an empty and elongated sheath between them (Figure 4 C arrows). In addition, the link between the oocyte and NCC units appeared to be elongated (Figure 4 C and K, arrows compared to F, arrow). As a proxy for successful maturation of a mature oocyte, we measured the proportion of NCCs dumping by measuring the volume difference between of the maximum and the last NCC. Overall, we found that the proportion of dumping was significantly decreased by caging (Wald χ^2^ = 18.86, *d*.*f*. = 4, p = 0.001; Figure 2 and S4). Interestingly, in EC caged queens, some nurse cell dumping occurred prematurely, just after the tipping point, when the oocyte was still underdeveloped (Figure 4 D arrowhead). As the days of EC caging progressed, the oocyte also appeared increasingly underdeveloped when nurse cell dumping occurred (Figure 4 E and F). By day seven of EC caging, some NCCs were at the oviduct end of the ovariole, but had not yet undergone PCD (Figure 1 J). In addition, there were some NCCs that had not collapsed even though the nurse cells had undergone PCD (Figure 4 B, F and G).

The oocytes closest to the oviduct undergo degeneration during the late vitellarium stage in EC caged queens. Initially, these degrading oocytes disconnect from their accompanying follicle cells (Figure 4 H arrow). As degradation progresses, the oocyte begins to collapse (Figure 4 I, white arrow) and undergo PCD (Figure 4 J arrow), until it degrades completely (see Figure 1 J). At the oviduct end of the ovariole, an actin aggregation accumulates as the days of EC caging increase (Figure 1G, J and M, arrow). In contrast, in the queens caged IC, the late vitellarium stage oocytes looked similar to those of the control and there appeared to be no actin aggregation (Figure 1 P).

In summary, confocal microscopy demonstrated that queens caged outside the colony (EC), had abnormal oogenesis as early as the fourth day post caging (Figure 1E, F, G). Throughout the days of EC caging the ovary area decreased, and the length of the germarium increased. This lengthening was a consequence of delayed growth of both the oocytes and NCC, as evidenced by the tipping point shifting towards the oviduct end. This latter shift resulted in underdeveloped, small, oocytes in the late vitellarium stage. By day seven post EC caging, only half of the NCCs completed ‘dumping’. In contrast, caging in the colony (IC) did not have such an extreme effect on queen ovaries. Although the size of the ovary slightly decreased as a result of caging IC, it is possible that this decrease was a result of the increase in the length of the germarium. However overall, queens caged IC retained healthy oocytes and may have continued some egg laying.

### Gene expression

The expression level of five (*Anarchy, Buffy, Tudor, Lncov1 and Draper*) of the six PCD-associated genes investigated increased with the duration of caging EC and IC (Wald χ^2^, p<0.05, table S2; Figure S5 A-E). The only exception to this pattern was *Ark*, where we found no effect of treatment on its expression levels (Wald χ^2^ = 5.43, *d*.*f* = 4, p = 0.246, table S2, Figure S5 F).

The expression of the four genes associated with nutrition sensitivity pathways and JH (*Kr-h1, Egfr, Tor* and *Foxo*) were significantly affected by the caging treatments (EC and IC) (Wald χ^2^, p<0.05, Figure S5 G-J, Table S2). These genes significantly increased in expression by day 10 of caging EC. The only gene that differed from this pattern was *Foxo*, which initially increased in expression before returning to levels similar to those seen in the control by day 10 of both EC and IC caging (Figure S5 J).

Gene expression of *Vg*, one of the two genes associated with vitellogenesis, was not affected by EC or IC caging (Wald χ^2^ = 4.44, *d.f* = 4, p = 0.35, Figure S5 K), and was extremely low. The expression of its receptor, *VgR*, was also not affected by caging treatments (Wald χ^2^ = 8.4, d.f = 4, p = 0.077, Figure S5 L).

## Discussion

Our study reveals that the control of oogenesis in honey bee queens uses different checkpoints to workers but similar genetic pathways. We identified three putative reproductive ‘checkpoints’ (also known as ‘control points’ in Ronai, Vergoz et al. 2016) that regulate oogenesis in honey bee queens. First, there is an absence of visible germline cells. This reduction in germline cells could either be due to suppression of stem cell differentiation (Figure 5 Checkpoint 1) or the early germline cell clusters are undergoing PCD (Figure 5 checkpoint 2).

**Figure 5.**
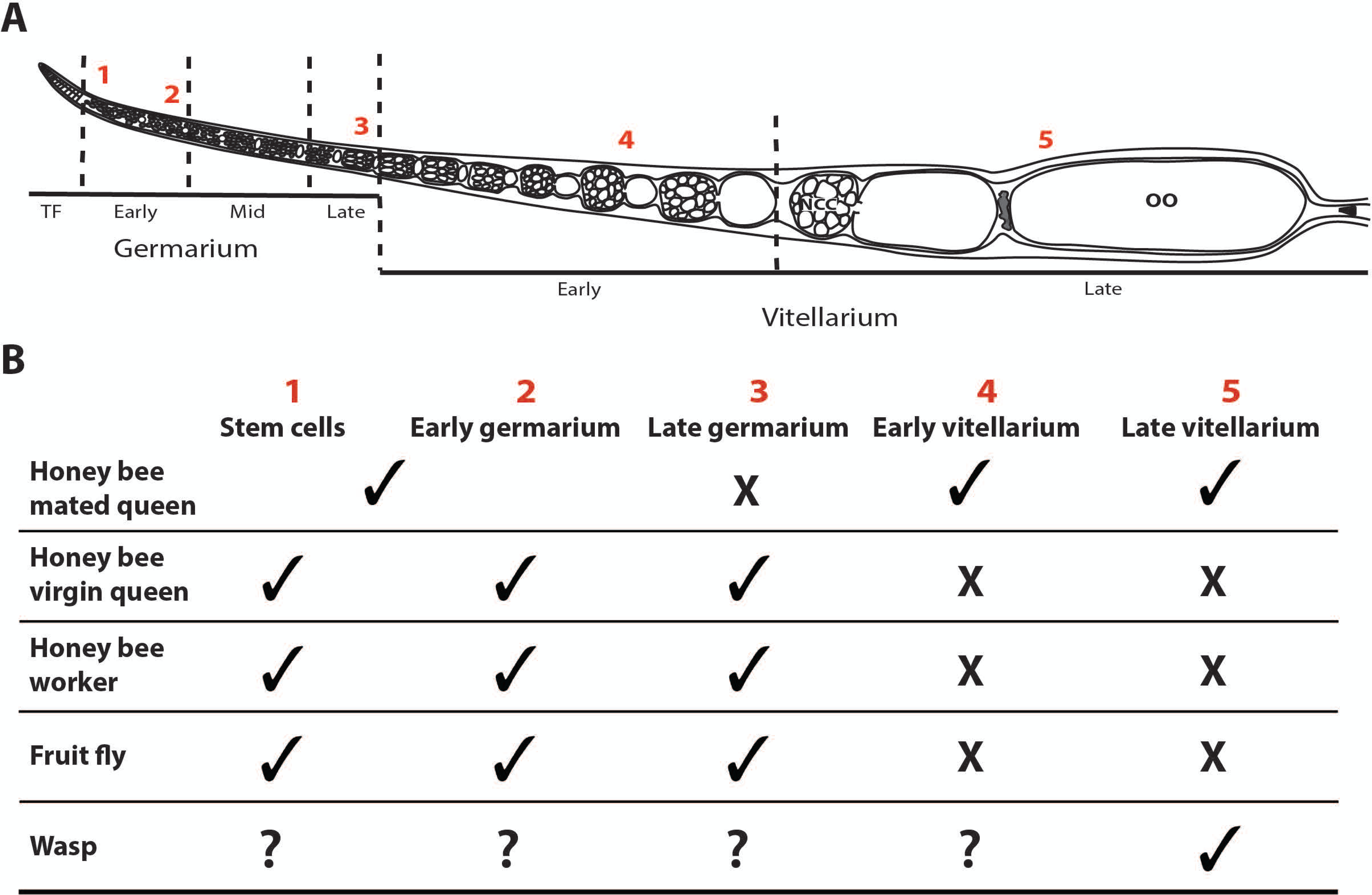
The oogenesis checkpoints in insects. (A) position of each checkpoint along an ovariole; (B) checkpoints used by the honey bee (*Apis mellifera*) mated queen, virgin queen, worker; solitary insect the fruit fly (*Drosophila melanogaster*); and another social insect the parasitoid wasp (*Nasonia vitripennis*).

PCD in the early germarium has previously been identified in virgin honey bee queens prevented from mating (Berger and Abdalla 2005; Cruz-Landim, Patrício et al. 2006) and in sterile honey bee workers (Tanaka, Schmidt-Capella et al. 2006; Ronai, Barton et al. 2015; Duncan, Hyink et al. 2016). These early checkpoints are also found in *Drosophila melanogaster* females that are under nutritional stress (Drummond-Barbosa and Spradling 2001; Jenkins, Timmons et al. 2013). Secondly, we found degradation of the oocytes in caged queens at the early vitellarium stage (Figure 1 I, L and 4 A, B). This checkpoint is not known to occur in any other hymenopteran species, or in insects in general, and is potentially unique to honey bee queens (Figure 5 checkpoint 4). Thirdly, mature oocytes degenerate in the late-vitellarium stage in honey bee queens prevented from laying (see also Cruz-Landim, Patrício et al. 2006) (Figure 1 G, J, M and 4 I, J). This checkpoint is also known to occur in another hymenopteran the parasitoid wasp (*Nasonia vitripennis*) (Hopkins and King 1964) but has not been shown in honey bee workers or other insects (Figure 5 checkpoint 5). Thus, it appears that honey bee queens have two additional checkpoints when compared to workers.

Our results indicate that honey bee queens and workers modulate oogenesis using a different combination of checkpoints to meet their different life history needs. In sterile workers the last checkpoint is at the late germarium stage (Tanaka and Hartfelder 2004; Ronai, Barton et al. 2015). This late germarium checkpoint is also found in nutritionally stressed *Drosophila* females (Drummond-Barbosa and Spradling 2001; Jenkins, Timmons et al. 2013) and other insects (Velentzas, Nezis et al. 2003; Uchida, Nishizuka et al. 2004; Mpakou, Velentzas et al. 2011). For honey bee workers, investing in oogenesis when there is no prospect of producing offspring is wasteful (Berger and da Cruz-Landim 2009). Therefore, it is advantageous for them to abort oocyte production before vitellogenesis, which is the most costly stage of oogenesis (Giorgi and Deri 1976; Drummond-Barbosa and Spradling 2001; Ronai, Vergoz et al. 2016). However, when the queen is absent or lost, there is clearly a fitness advantage for workers that are able to quickly switch on oogenesis (Page and Robinson 1994). In contrast to workers, as shown in our study, queens use a combination of checkpoints that act at later stages of oocyte development. These later checkpoints allow the queen to reabsorb the oocytes and therefore preserve invaluable resources if the egg cannot be laid. Therefore, queens slow down the rate of oocyte production while continuing to produce oocytes.

PCD is an important mechanism for modulating oogenesis in insects and our morphological results support this idea by showing clear evidence of PCD in the ovarioles of caged honey bee queens. Therefore, it was unsurprising to find direct evidence of cell death indicated by the higher expression levels of the phagocytosis engulfment receptor gene *Draper* (Etchegaray, Timmons et al. 2012) in caged queens. Furthermore, we have shown an upregulation of genes from two independent PCD pathways in honey bee queens: one that involves *Anarchy* and *Buffy*, a mitochondrial regulatory gene network associated with PCD (Tanner, Blute et al. 2011) and another, the Lncov1/Tudor-SN complex, which interferes with effector caspase activity (Hartfelder, Tiberio et al. 2018). Interestingly, the expression pattern of *Anarchy* and *Buffy* in caged queens is not the same as in virgin queens with non-activated ovaries (Ronai, Oldroyd et al. 2016). Rather, their expression patterns are correlated with the suppression of oogenesis in workers exposed to the queen (Ronai, Oldroyd et al. 2016). Interestingly, expression of *Ark*, a pro-apoptotic gene that initiates the caspase cascade resulting in PCD (Cain, Bratton et al. 2002; Dallacqua and Bitondi 2014), remained constant across our treatments. In virgin queens with non-activated ovaries, *Ark* is downregulated but is not differentially expressed between sterile and laying workers (Rodriguez, Oliver et al. 1999; Dallacqua and Bitondi 2014; Ronai, Oldroyd et al. 2016). Thus, our results suggest that under stressful environmental conditions queens modulate their oogenesis using the same mechanism (PCD) as workers and do not revert to a virgin-like status.

A critical process for the maturation of an oocyte in insects is the accumulation of vitellogenin, the main egg yolk protein (Raikhel and Dhadialla 1992; Tufail and Takeda 2008). Vitellogenin is mostly synthesised in the fat body and then transferred to the oocyte via the vitellogenin receptor (*VgR*) (Engels 1974). Therefore, it was not surprising that *Vg* expression in the ovary was extremely low (Guidugli, Nascimento et al. 2005; Cardoso‐Júnior, Oldroyd et al. 2021) and did not change as a result of the treatments. However, previous studies have used *VgR* expression levels as a proxy for the reproductive state of honey bee workers (Guidugli-Lazzarini, do Nascimento et al. 2008; Cardoso‐Júnior, Oldroyd et al. 2021), therefore one may expect *VgR* expression levels to decrease as a result of the reduced oogenesis in caged queens. Our results suggest that under stressful conditions queens potentially keep *VgR* levels constant in the ovaries in order to be able to swiftly provide vitellogenin and restore oogenesis when the environmental conditions improve.

Our results emphasise the significance of the queens’ environment on her reproductive capacity and suggest that nutrition and a normal social environment are necessary for regular oogenesis. We observed a distinct difference in the effects of caging, between EC and IC on oogenesis. Queens caged within the colony had healthier ovaries and oocytes than externally caged queens, in which oogenesis was more severely disrupted. This difference may be linked to the amount of care provided by the workers, particularly the quantity and quality of food fed by the escort workers to the queens EC relative to that which could be provided IC (Szabo 1974). As is standard beekeeping practise, the only source of nutrition available to the escort workers in our external cages was ‘queen candy’ (Laidlaw and Page 1997). While candy is rich in carbohydrates, it lacks fat and protein, the latter being essential for oogenesis (Fine, Shpigler et al. 2018). Thus, one would expect the ability of escort workers to feed the queen in their care with glandular secretions is severely curtailed because the workers themselves are nutritionally stressed.

Caging queens both EC and IC influenced the expression of genes associated with the nutrient sensitive pathways Tor and EGFR in our study. We suggest that the upregulation of the nutrition-sensitive genes in the caged and banked queens was due to nutritional stress. *FoxO*, a downstream target gene of *Tor* and *Egfr*, is also upregulated in nutritionally-stressed honey bee workers (Wheeler, Buck et al. 2014; Cardoso‐Júnior, Oldroyd et al. 2021) and in nutritionally-stressed solitary insects such as *Drosophila* (Jouandin, Ghiglione et al. 2014). However, Tor has been shown to be downregulated in ovaries of nutritionally-stressed Drosophila (Pritchett and McCall 2012; Smykal and Raikhel 2015) and mosquitoes (Hansen, Attardo et al. 2005; Perez-Hedo, Rivera-Perez et al. 2014). The up-regulation of Kr-h1, a proxy for juvenile hormone (JH) titres (Minakuchi, Namiki et al. 2009; Belles and Santos 2014; Lago, Humann et al. 2016) observed in the queens caged both EC and IC is consistent with higher levels of JH seen in non-reproductive honey bee workers (Robinson, Strambi et al. 1992) and in nutritionally stressed *Drosophila* (Terashima and Bownes 2004) Therefore, JH possibly works as an intermediate regulator of adult ovary activation in both honey bee castes.

Although more studies need to be conducted to assess the effect of nutrition quantity and quality on honey bee queen oogenesis, our study suggests that nutrition could serve as a cue for the suppression of oogenesis in queens. Our findings accord with behavioural observations (Allen 1956; Pierce, Lewis et al. 2007) in which queens are fed less during swarming. In our particular study, IC queens acquired their nutrition via trophalaxis from colony workers through their cage. Food obtained from free-running nurse bees would be almost certainly of higher quality to that available to the EC queens. Whilst nutrition is not the only environmental difference between queens cages EC and IC, we suggest that food is a major factor that contributed to the nearly-normal ovary morphology of the IC queens, and the loss of oocytes in the EC queens. Nutrition also regulates oogenesis in solitary insects such as *Drosophila* (Drummond-Barbosa and Spradling 2001; Gruntenko, Bownes et al. 2003; Barth, Szabad et al. 2011).

Our study has practical ramifications for beekeepers. It is clear that queens caged in the colony fare better than queens caged externally to the colony, and that keeping queens within the colony environment should be the preferred method for queen storage. It is suggested that the period of caging should be minimised. Likely, reformulating the candy diet for caged queens and their escorts to include pollen as a source of protein and fat would enhance queen welfare in the mail, and we recommend that this be investigated in a future study.

In conclusion, although queens and worker honey bees share the same regulatory template, they have diverged in how they regulate oogenesis. Reproductive division of labour between castes is fundamental for the evolution of eusociality in the social insects. One explanation of the creation of castes is the “reproductive ground plan hypothesis” (West-Eberhard 1987) which suggests that the queen and worker castes evolved from a decoupling of the reproductive and foraging stages of a solitary ancestral insect from which contemporary eusocial bees are derived (Amdam, Norberg et al. 2004). This decoupling resulted in the molecular, morphological and behavioural differences between the queen and worker castes but preserved the underling pathways that regulate nutrition and reproduction. It is proposed here that the reproductive divergence of queen and worker honey bees is achieved through different oogenesis checkpoints but the same PCD pathways in order to modulate oogenesis and to meet the reproductive divergence of the two castes.

## Acknowledgements

To Rhiarn Hoban for drawing the detailed ovariole structure; to Mackenzie R. Lovegrove for her assistance in trouble shooting the fixation stages of the ovarioles and to the members the Behaviour and Genetics of Social Insects Lab for their assistance and advice.

## Conflict of interests

We have no conflict of interests.

## Data archiving

All supporting datasets for this paper are available as part of the electronic supplementary material.

## Funding

This work was funded by the Australian Research Council project DP180101696.

---

Figure S1

Number of visible oocytes along an ovariole of honey gee queens from the control, caged in-colony (IC) (4, 7 or 10 days) or caged Ex-colony (EC) (10 days) treatment groups. n = 4 control, caged IC 10 days and caged EC 10 days; n = 5 caged EC 4 and 7 days. Error bars are SE of the means.

Figure S2

Spherical radius (*R*_eq_) (μm) of oocytes and NCC of honey bee queens from the control, caged in-colony (IC) (4, 7 or 10 days) or caged Ex-colony (EC) (10 days) treatment groups. (A) maximum oocyte radius; (B) maximum nurse cell chamber radius; (C) minimum oocyte radius; (D) minimum nurse chamber radius. Sample size see Figure S1. Error bars are SE of the means. Bars with a different letter are significantly different (*p* < 0.05).

Figure S3

Position of the oocyte in the ovariole when the tipping point occurs. Honey bee queens from the control, caged in-colony (IC) (4, 7 or 10 days) or caged Ex-colony (EC) (10 days) treatment groups. Sample size see Figure S1. Error bars are SE of the means. Bars with a different letter are significantly different (*p* < 0.05).

Figure S4

Extent of nurse cell dumping in honey bee queens from the control, caged in-colony (IC) (4, 7 or 10 days) or caged Ex-colony (EC) (10 days) treatment groups. Sample size see Figure S1. Error bars are SE of the means. Bars with a different letter are significantly different (*p* < 0.05).

Figure S5

Gene expression in the ovaries of honey bee queens from the control, caged in-colony (IC) (4, 7 or 10 days) or caged Ex-colony (EC) (10 days) treatment groups. Genes related to program cell death (A) *Anarchy* (B) *Lncov-1* (C) *Tudor* (D) *Buffy* (E) *Draper* and (F) *Ark*. Genes related to nutrition (G) *Kr-h1*; (H) *Egfr*; (I) Tor and (J) *FoxO*. Genes related to vitellogenesis (K) *Vg* and (L) *VgR*. Sample size see Figure S1. Boxplot whiskers represent minimum and maximum values, the box is defined by 25th percentile, median and 75th percentile. Bars with a different letter are significantly different (*p* < 0.05).

## Notes

### Competing Interest Statement

The authors have declared no competing interest.

### Summary of Updates

A mistake in one of the authors names and the order of the figures

