## Supplementary figures 1-5 for "Regulation of oogenesis in the queen honey bee (*Apis mellifera*)"

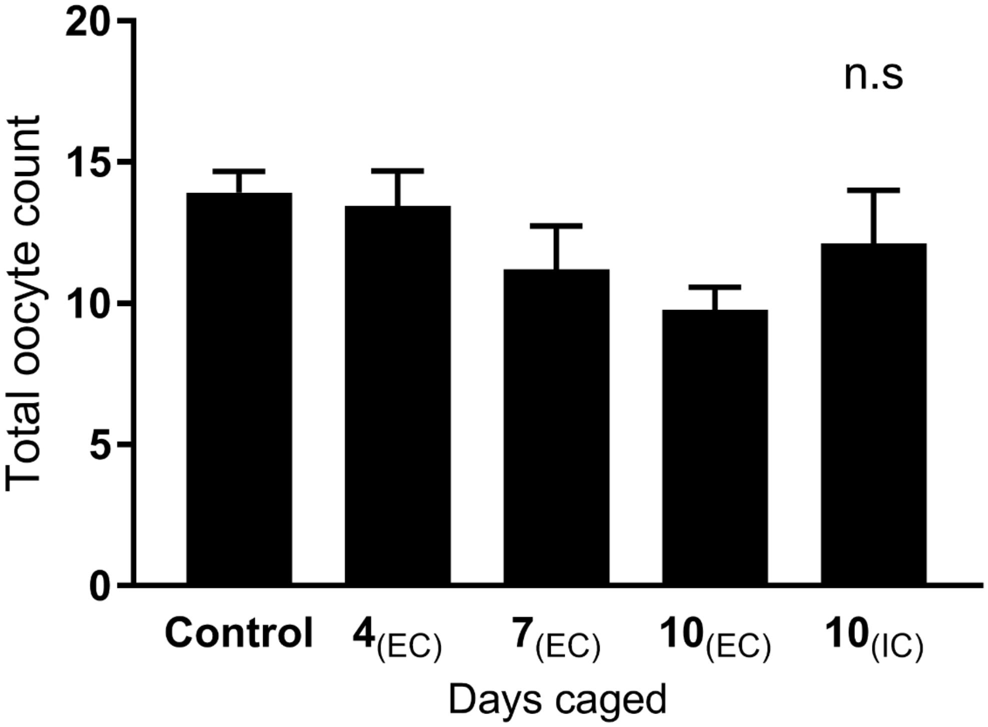


**Figure S1.** Number of visible oocytes along an ovariole of honey gee queens from the control, caged in-colony (IC) (4, 7 or 10 days) or caged Ex-colony (EC) (10 days) treatment groups. n = 4 control, caged IC 10 days and caged EC 10 days; n = 5 caged EC 4 and 7 days. Error bars are SE of the means.

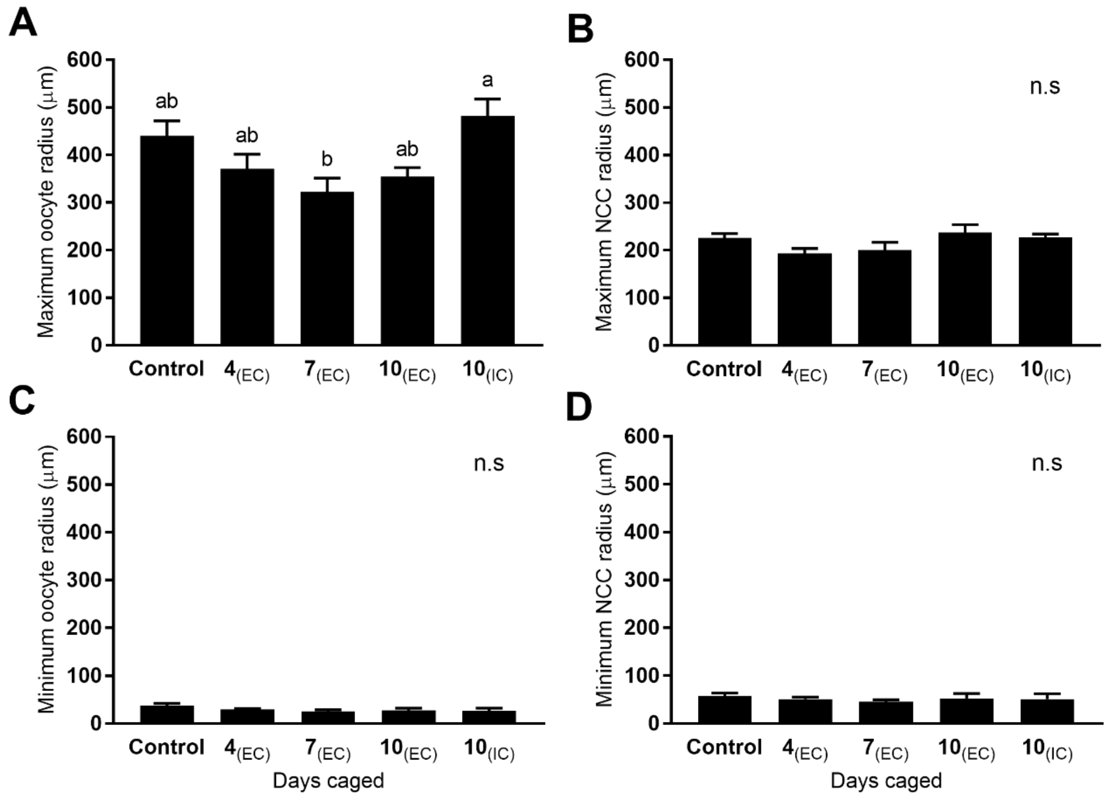


**Figure S2.** Spherical radius ($R_{eq}$) (µm) of oocytes and NCC of honey bee queens from the control, caged in-colony (IC) (4, 7 or 10 days) or caged Ex-colony (EC) (10 days) treatment groups. (A) maximum oocyte radius; (B) maximum nurse cell chamber radius; (C) minimum oocyte radius; (D) minimum nurse chamber radius. Sample size see Figure S1. Error bars are SE of the means. Bars with a different letter are significantly different ($p$ < 0.05).

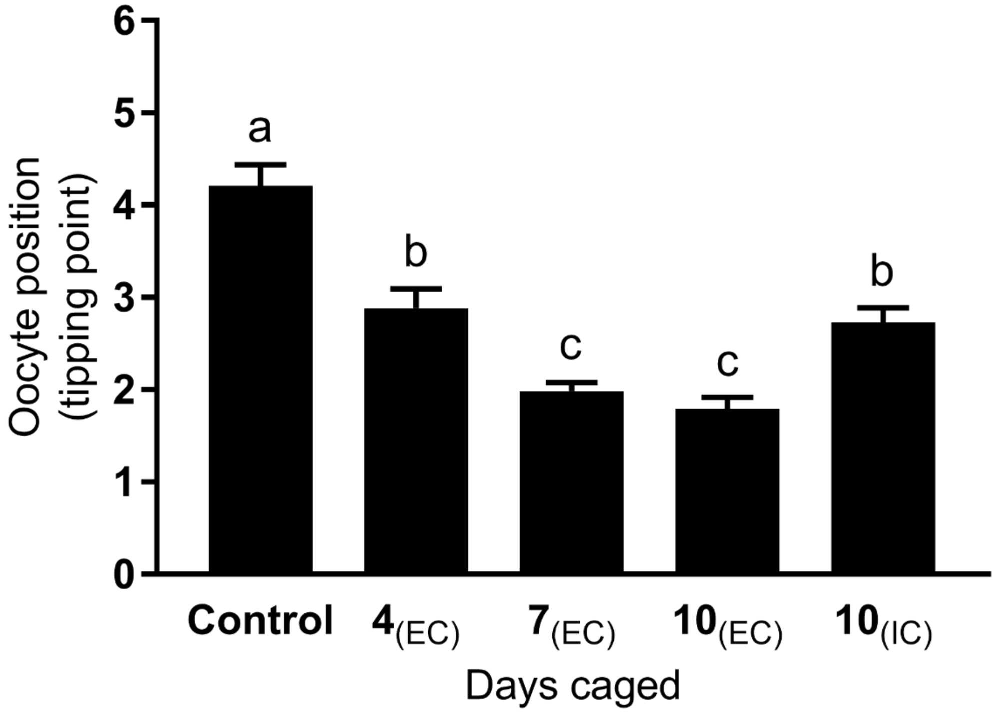


**Figure S3.** Position of the oocyte in the ovariole when the tipping point occurs. Honey bee queens from the control, caged in-colony (IC) (4, 7 or 10 days) or caged Ex-colony (EC) (10 days) treatment groups. Sample size see Figure S1. Error bars are SE of the means. Bars with a different letter are significantly different ($p < 0.05$).

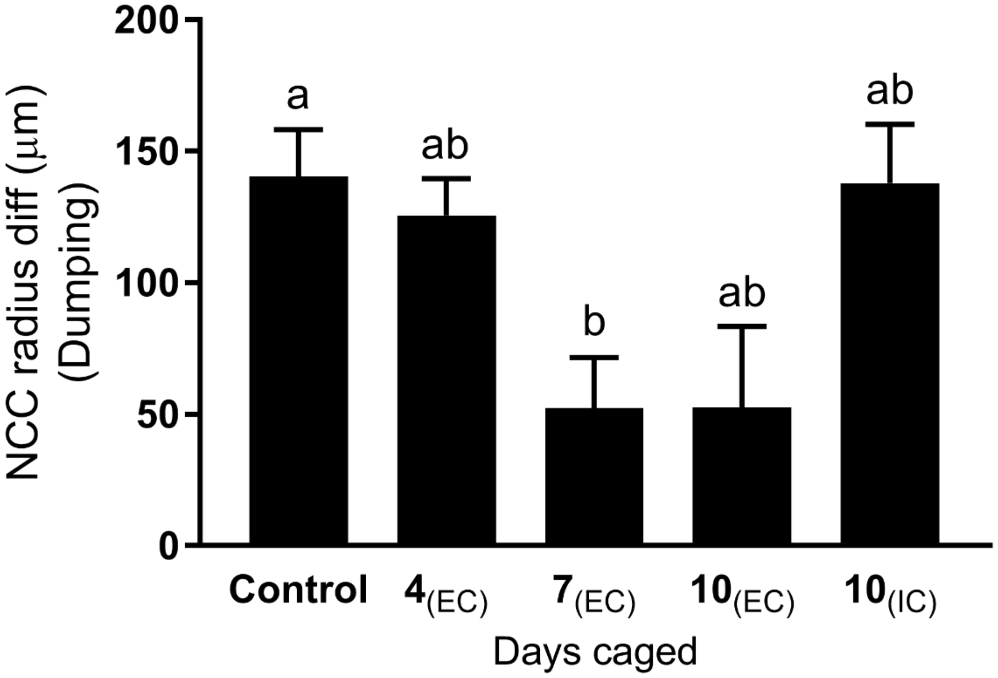


**Figure S4.** Extent of nurse cell dumping in honey bee queens from the control, caged in-colony (IC) (4, 7 or 10 days) or caged Ex-colony (EC) (10 days) treatment groups. Sample size see Figure S1. Error bars are SE of the means. Bars with a different letter are significantly different ($p$ < 0.05).

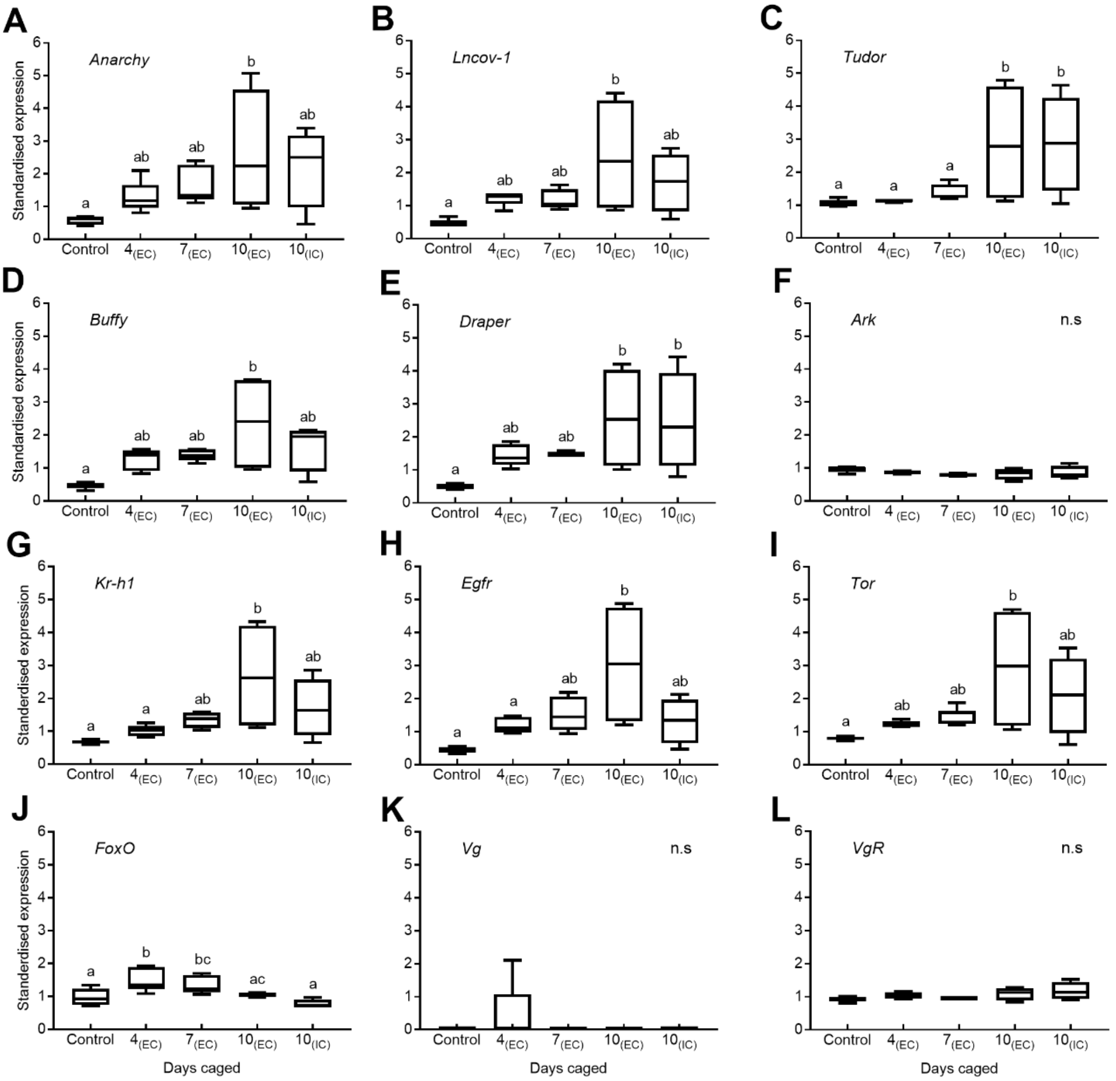


**Figure S5.** Gene expression in the ovaries of honey bee queens from the control, caged in-colony (IC) (4, 7 or 10 days) or caged Ex-colony (EC) (10 days) treatment groups. Genes related to program cell death (A) *Anarchy* (B) *Lncov-1* (C) *Tudor* (D) *Buffy* (E) *Draper* and (F) *Ark*. Genes related to nutrition (G) *Kr-h1*; (H) *Egfr*; (I) *Tor* and (J) *FoxO*. Genes related to vitellogenesis (K) *Vg* and (L) *VgR*. Sample size see Figure S1. Boxplot whiskers represent minimum and maximum values, the box is defined by 25th percentile, median and 75th percentile. Bars with a different letter are significantly different ($p$ < 0.05).
