## Supplementary methods S1 for "Regulation of oogenesis in the queen honey bee (*Apis mellifera*)"

#### Supplementary method Image processing

##### Annotating the cells

1. A maximum intensity projection of the Z-stack was performed.
2. A gamma transformation (typically 0.6) was applied so the less intense regions of the image could be visualised
3. Starting with the largest cells in the strand, each pair of nurse and oocyte were annotated using the ‘freehand tool’ and added to the Region Of Interest (ROI) Manger.

The ROI lists were labelled and sorted into chains and unit type, starting from the oocyte ‘o’ followed by the nurse cell chamber (NNC) ‘n’ interchanging down the chain until the borders between the two units where no longer clear (Figure 1).

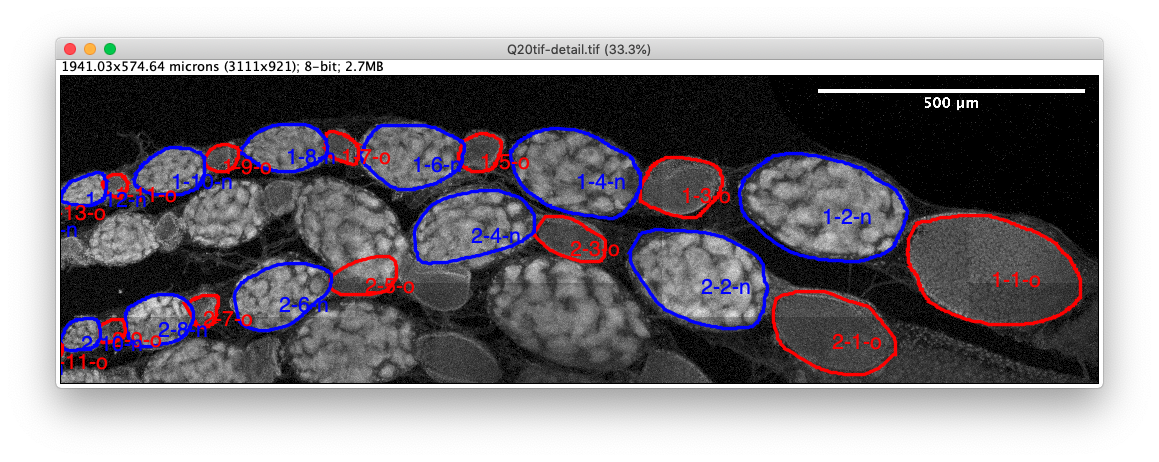

Figure 1: example of labeled annotation for 2 ovarioles. Always starting from the largest oocyte such as the odd numbers are oocytes and even numbers are NCC. Screen shot from ImageJ.

#### Volume estimate

The volume of each unit cell ROI was calculated as following:

- The maximum and minimum Feret’s diameterwere measured
- The oblate volume was approximated using the measured maximum and minimum Feret’s diametrer as follows$V_{o}= \frac{4}{3}\times\pi\times maxFeret\times{minFeret}^{2}$
- The equivalent spherical radius, $R_{\text{eq}}$, was calculated used in subsequent statistical analyses:

$$R_{eq}=\sqrt[3]{\frac{3}{4\pi}V_{o}}$$

### Aliening of the units

As the cell units (oocyte and nurse) traverse along the ovary they change in size (Figure 3a). It is convenient to use the effective spherical radius, $R_{eq}$ (herby ‘radius’), as the volume changes over a large range (Figure 3b). This is equivalent to a cube root transformation of the data which is common in statistical analysis. Initially (at large cell positions) the nurse cells are larger than the oocytes. As the cells grow (cell position decreases) the oocytes grow faster than the nurse cells, and the oocyte cells eventually overtake the nurse cells. Before the oocyte leaves the ovary the nurse cell may undergo a rapid decrease in size, a process that is referred to as dumping. This usually occurs within 1-3 cell positions from the end (Figure 3c).

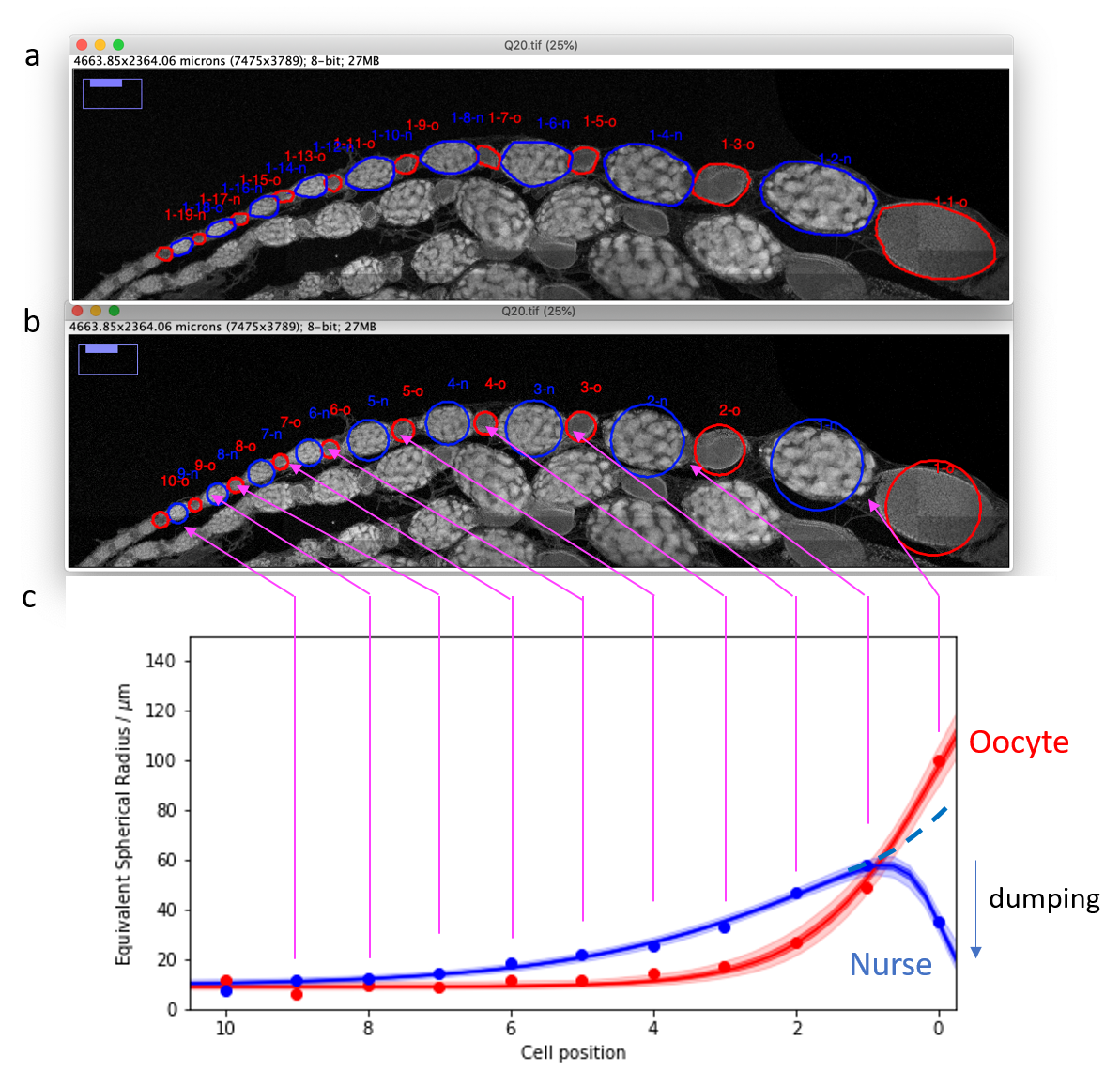

Figure 3: Work flow for the analysis of the ovariole. Oocyte (red) NCC (blue). a) Manually annotated ovariole; b) Conversion to equivalent spherical radius, $R_{\text{eq}}$; c) Schematic describing the changes in each of the cell units along the ovariole. Screen shot from ImageJ.

##### Parameter used for the statistical analysis

1) Maximum and minimum volume of the oocyte and NNC.
2) Tipping point: The cell position where the oocyte and NNC are most similar in volume.
3) Dumping: The volume change between the maximum and first NNC.
