## Supplementary table 1-2 for "Regulation of oogenesis in the queen honey bee (*Apis mellifera*)"

Supplementary table S1: Primer list used for RT-qPCR.

| Gene | Sequence 5’-3’ | GenBank | Reference | Efficiency |
| --- | --- | --- | --- | --- |
| *Anarchy* | F: ACAAAGGAATGGAAGCCAAA | GB48961 | (Ronai, Oldroyd et al. 2016) | 1.78 |
|  | R: GGAAATTTACACGGGGAACA |  |  |  |
| *Buffy* | F: TGTTGCACGACAAAT | GB49154 | (Ronai, Oldroyd et al. 2016) | 1.74 |
|  | R: AGGTTTGCCCTGACG |  |  |  |
| *VgR* | F: ACCTTACGACATTGCCCT | GB40823 | (Lourenço, Martins et al. 2012) | 1.72 |
|  | R: TGTGATTTTCGGTCCAAGCCC |  |  |  |
| *Actin* | F: TGCCAACACTGTCCTTTCTG | GB44311 | (Scharlaken, de Graaf et al. 2008) | 2.03 |
|  | R: AGAATTGACCCACCAATCCA |  |  |  |
| *pRS5* | F: AATTATTTGGTCGCTGGAATTG | GB45730 | (Ronai, Oldroyd et al. 2016) | 1.68 |
|  | R: TAACGTCCAGCAGAATGTGGTA |  |  |  |
| *S6k* | F: GCTTACTGGAATGCC | GB45876 | (Ronai, Oldroyd et al. 2016) | 1.69 |
|  | R: GGGTCCAGATCCCAA |  |  |  |
| *Kr-h1* | F: GCACTGGCAGTGACAAGGAA | GB45427 | (Fussnecker and Grozinger 2008) | 1.73 |
|  | R: CGTGGAGTGTTATCGTAAGTAGCAA |  |  |  |
| *Tudor* | F: GTTACCAGTGGATCGCGTCT | GB40977 | (Tiberio, et al. 2017) | 1.67 |
|  | R: AGCAATGCTCTCGGGTGAAA |  |  |  |
| *FoxO* | F: TTATGCGAGTGCAGAACGAG | GB48301 | (Wheeler, Buck et al. 2014) | 1.81 |
|  | R: AAGCGGACTGTCTGGAAAGA |  |  |  |
| Lncov1 | F: GGAGAAGCTTTGGGGAGAG | (intron 5 of LOC726407 – GB45056) | (Humann and Hartfelder 2011) | 1.77 |
|  | R: CTGCTACACACCACCATAAC |  |  |  |
| Tor | F: AACAACTGTTGCTGACGGTG | GB44905 | (Patel, Fondrk et al. 2007) | 1.67 |
|  | R: GTTGCAGTCCAGGCTTTTTG |  |  |  |
| EGFR | F: GAGATCCTGGCCGATATCTTG | GB54477 | (Marco Antonio and Hartfelder 2017) | 1.77 |
|  | R: GCAGAAAGTCCTGGTGGAATC |  |  |  |
| Draper | F: CAGCAGGATGGGTTGGTG | GB54664 | This Study | 1.73 |
|  | R: CCATTTTGTGGATTGCAGGC |  |  |  |
| Vg | F: CCGACGAGGACCTGTTGATTA | GB49544 | (Koywiwattrakul, Thompson et al. 2005) | 1.88 |
|  | R: CTAGGATACGTGGTCATGACA |  |  |  |
| Ark | F: AGTTGTGGTCCAGGT | GB52453 | (Ronai, Oldroyd et al. 2016) | 1.7 |
|  | R: GTTGGAGGAAGAGTGGGTGT |  |  |  |
| Ef1a | F: TGCAACCTACTAAGCCGATG | GB52028 | (Ronai, Oldroyd et al. 2016) | 1.74 |
|  | R: GACCTTGCCCTGGGTATCTT |  |  |  |
| RF49 | F: CGTCATATGTTGCCAACTGGT | GB47227 | (Lourenço, Mackert et al. 2008) | 1.69 |
|  | R: TTGAGCACGTTCAACAATGG |  |  |  |

Supplementary table S2: GLMM test for gene expression analysis. Dependent variable: expression level, Model: (Intercept), Time.

| **Wald** |  |  |  |  |
| --- | --- | --- | --- | --- |
| Group | Chisq | Df | Pr(>Chisq) | Sig |
| Ark | 5.43 | 4 | 0.246 |  |
| foxo | 25.05 | 4 | 0.000 | *** |
| Vg | 3.47 | 4 | 0.482 |  |
| VgR | 8.43 | 4 | 0.077 | . |
| Anarchy | 11.83 | 4 | 0.019 | * |
| Buffy | 17.11 | 4 | 0.002 | ** |
| Lancov1 | 14.34 | 4 | 0.006 | ** |
| Tudor | 15.86 | 4 | 0.003 | ** |
| Kr-h1 | 16.61 | 4 | 0.002 | ** |
| Tor | 14.45 | 4 | 0.006 | ** |
| EFGR | 21.00 | 4 | 0.000 | *** |
| Draper | 16.13 | 4 | 0.003 | ** |
| Signif. codes: 0 ‘***’ 0.001 ‘**’ 0.01 ‘*’ 0.05 ‘.’ 0.1 ‘ ’ 1 | | | | |

| **Wald** |  |  |  |  |
| --- | --- | --- | --- | --- |
| Group | Chisq | Df | Pr(>Chisq) | Sig |
| Max nc | 8.97 | 4 | 0.062 | . |
| Max o | 18.79 | 4 | 0.001 | *** |
| Min nc | 1.48 | 4 | 0.830 |  |
| Min o | 5.43 | 4 | 0.246 |  |
| Tipping | 119.43 | 4 | 0.000 | *** |
| Dumping | 18.86 | 4 | 0.001 | *** |
| Length | 6.04 | 4 | 0.196 |  |
| Ovary area | 96.25 | 4 | 0.000 | *** |
| Signif. codes: 0 ‘***’ 0.001 ‘**’ 0.01 ‘*’ 0.05 ‘.’ 0.1 ‘ ’ 1 | | | | |

Dallacqua R.P., M.M.G. Bitondi. (2014) Dimorphic ovary differentiation in honeybee (*Apis mellifera*) larvae involves caste-specific expression of homologs of Ark and Buffy cell death genes. PloS one **9**(5): e98088.

Fussnecker B., C. Grozinger. (2008) Dissecting the role of Kr‐h1 brain gene expression in foraging behavior in honey bees (*Apis mellifera*). Insect Mol. Biol. **17**(5): 515-522.

Humann F.C., K. Hartfelder. (2011) Representational Difference Analysis (RDA) reveals differential expression of conserved as well as novel genes during caste-specific development of the honey bee (*Apis mellifera* L.) ovary. Insect Biochem. Mol. Biol. **41**(8): 602-612.

Koywiwattrakul P., G.J. Thompson, S. Sitthipraneed, B.P. Oldroyd, R. Maleszka. (2005) Effects of carbon dioxide narcosis on ovary activation and gene expression in worker honeybees, *Apis mellifera*. J Insect Sci **5**(1): 36.

Lourenço A.P., A. Mackert, A. dos Santos Cristino, Z.L.P. Simões. (2008) Validation of reference genes for gene expression studies in the honey bee, *Apis mellifera*, by quantitative real-time RT-PCR. Apidologie **39**(3): 372-385.

Lourenço A.P., J.R. Martins, K.R. Guidugli-Lazzarini, L.M.F. Macedo, M.M.G. Bitondi, et al. (2012) Potential costs of bacterial infection on storage protein gene expression and reproduction in queenless *Apis mellifera* worker bees on distinct dietary regimes. J. Insect Physiol. **58**(9): 1217-1225.

Marco Antonio D.S., K. Hartfelder. (2017) Toward an understanding of divergent compound eye development in drones and workers of the honeybee (Apis mellifera L.): a correlative analysis of morphology and gene expression. Journal of Experimental Zoology Part B: Molecular and Developmental Evolution **328**(1-2): 139-156.

Patel A., M.K. Fondrk, O. Kaftanoglu, C. Emore, G. Hunt, et al. (2007) The making of a queen: TOR pathway is a key player in diphenic caste development. PloS one **2**(6): e509.

Ronai I., B.P. Oldroyd, D.A. Barton, G. Cabanes, J. Lim, et al. (2016) Anarchy is a molecular signature of worker sterility in the honey bee. Mol. Biol. Evol. **33**(1): 134-142.

Scharlaken B., D.C. de Graaf, K. Goossens, M. Brunain, L.J. Peelman, et al. (2008) Reference gene selection for insect expression studies using quantitative real-time PCR: The head of the honeybee, Apis mellifera, after a bacterial challenge. J Insect Sci **8**(1): 33.

Wheeler D., N. Buck, J. Evans. (2014) Expression of insulin/insulin‐like signalling and TOR pathway genes in honey bee caste determination. Insect Mol. Biol. **23**(1): 113-121.
